# New rice varieties with improved phosphorus-efficiency for low-input smallholder rice production in Africa

**DOI:** 10.64898/2025.12.06.692696

**Authors:** M Wissuwa, HN Ranaivo, MT Rakotondramanana, S Rafaliarivony, K Kondo, Y Ueda, LT Dinh, M Connor, JH Chin, J Pariasca Tanaka

**Author notes:** Corresponding author: Matthias Wissuwa.

## Abstract

Smallholder farmers in Sub-Saharan Africa frequently produce rice in marginal environments where low soil fertility and other biotic and abiotic stresses limit productivity. Rice varieties developed by centralized breeding under favorable conditions on research stations have often not been adopted by farmers in such marginal environments. Our objective was to develop modern rice varieties adapted to such low-input conditions through combining pre-breeding research with subsequent selection and variety testing directly in the target environment: smallholder farmers’ fields in Madagascar with phosphorus (P) fixing soils.

Two breeding populations were developed for this purpose, one based on marker-assisted introgression of the Pup1 locus into IR64, the second using a donor (DJ123) for internal P utilization efficiency and external P acquisition efficiency. Selection within these populations was conducted in fields managed according to local farmers’ practice without mineral fertilizer addition. Selected breeding lines underwent government-supervised variety release testing including farmer participatory evaluations in four Malagasy regions between 18–1350 masl altitude, and two lines were released as varieties ’FyVary32’ and ‘VyVary85’. Both had between 0.47–0.8 t ha-1 higher grain yield than parent IR64 and local check X265. Yield advantages were stable across a range from 2.1–5.5 t ha-1 (national average: 2.8 t ha-1). Higher yields were accompanied by superior root development and P uptake and by more efficient internal P utilization in FyVary85. Results show that decentralized breeding in marginal environments can produce varieties not only superior in lowest-yielding environments but across a broader range, clearly surpassing national average yields.

## Introduction

Rice consumption in Sub-Saharan Africa (SSA) is increasing faster than local rice production, forcing many countries to increase rice imports from Asia (Otsuka et al. 2024; Yuan et al. 2024). One reason for this widening gap between local demand and supply are the relatively stagnant yields in SSA, which average 2.2 t ha^-1^ in West-Africa and 2.72 t ha^-1^ in East-Africa compared to yields in excess of 4.2 t ha^-1^ in tropical Asia (FAOSTAT, 2020). Where increases in total production per region were seen in SSA, this was more often a result of expanding land area cropped with rice rather than due to an increase in productivity (Yuan et al. 2024).

Low rice yields in SSA are caused by the combination of a high proportion of rainfed rice in combination with low soil fertility and insufficient fertilizer application (Haefele et al. 2014; Saito et al. 2019). Highly weathered infertile soils (Ferralsols, Lixisols) are prevalent in the region and their capacity to bind phosphorous (P) in forms of low plant availability frequently results in P being the most growth-limiting nutrient in rice production in SSA (Saito et al. 2019; Rakotoson et al. 2022). Despite efforts to increase fertilizer application rates through farmer education or government subsidies, average applications rates continue to lag far behind those seen in Asia (FAOSTAT, 2020; Rakotoson et al. 2022).

Madagascar is the 2^nd^ biggest rice producer in SSA after Nigeria, with a long tradition of rice cultivation and slightly above-average rice yields (2.87 t ha^-1^) for the region (FAOSTAT, 2020). Most of the rice is produced by smallholder farmers. Yield improvements from 2.8 t ha^-1^ to 4.3 – 4.8 t ha^-1^ were reported when farmers adopted a package including fertilizers, improved irrigation and high-quality certified seeds (JICA, 2021), indicating that yields comparable to tropical regions elsewhere can be achieved in Madagascar and possibly throughout SSA when the entire green-revolution package – irrigation, high fertilizer inputs, improved seeds, reduction of pests and diseases through application of agro-chemicals - is available (Otsuka et al. 2023). However, with 6.9 kg ha^-1^ fertilizer application in Madagascar, rates have remained stagnant and even lower than the average for SSA (18.2 kg ha^-1^) and far below the 185 kg ha^-1^ seen in South Asia (all data for 2022, World bank). This extremely low fertilizer use persists despite standard fertilizer recommendations in the range of 200-300 kg ha^-1^ of NPK (11-22-16) fertilizers being in existence for decades. However, chemical fertilizer are often inaccessible and costly for smallholder farmers (Vanlauwe et al., 2014). In Madagascar, one kg of diammonium phosphate (DAP) fertilizer costs the equivalent of 1.37 € in Malagasy village markets (Ranaivo, pers. comm.) compared to about 0.63-0.67 € in Europe and only 0.30 € for subsidized DAP in India. Furthermore, farmers’ education and their perception of viable returns on fertilizer use (Chianu et al., 2012) may further hinder widespread fertilizer application.

In the absence of encouraging country-wide trends in the adoption of green revolution rice production packages in Madagascar and much of SSA, alternative options for a sustainable intensification need to be pursued. These should be tailored to smallholder farmers and may include micro-dosing options (Rakotoson et al. 2020; Vandamme et al. 2016), ideally in combination with improved modern varieties adapted to prevailing conditions on these farms (Tsujimoto 2025). Vandamme et al. (2015) highlighted that a cost-efficient partial solution to the soil fertility problem in SSA would be the development of varieties with improved P acquisition and utilization efficiencies. This possible solution shall be explored for the case of rice variety development for smallholder farmers in Madagascar.

The conventional on-station breeding approach that seeks to select breeding lines with high yield potential under “ideal” high-input conditions may no longer be the best solution to achieve that goal. The prevalence of traditional rice varieties throughout Madagascar (Minten and Barrett, 2008) is an indication that plant breeding has not properly addressed the needs of the mostly resource-poor smallholder farmers. It is furthermore indicative of specific adaptations to lower soil fertility being present in such traditional varieties (Dröge et al., 2022). Under nutrient-limited conditions plant growth and ultimately grain yield will depend on the quantity of the limiting nutrient being taken up and for a nutrient with low mobility in the soil such as P, that will be a function of the soil volume explored by the root system, which is directly related to root system size (Mori et al. 2016), and the P acquisition efficiency (PAE) which relates to nutrient uptake per unit size (Mori et al. 2016). In addition, the internal P utilization efficiency (PUE) is key as it determines how much biomass is produced per unit nutrient taken up (Wissuwa et al. 2015).

Efforts have been undertaken to identify possible donors and underlying genetic factors for above P-efficiency traits. Possibly the most well-known locus enhancing P uptake in rice is the *Pup1* locus from donor accession ‘Kasalath’ and the corresponding causal gene *OsPSTOL1* that leads to better soil exploration through enhanced crown root development (Gamuyao et al. 2012). With *OsPSTOL1* having been identified through map-based cloning, sets of markers are available (Chin et al. 2011) to introgress this locus into different recipient varieties through marker assisted selection (MAS).

For other target traits, loci with comparable strength or precision of mapping are not available. Screening association panels of gene-bank accessions had identified large variation for root size (Mori et al. 2016) and smaller variation for PAE (Mori et al. 2016) or for PUE (Wissuwa et al. 2015). Each trait was apparently controlled by multiple small-effect QTL that have so far not been linked to a single underlying gene. However, several donors have been identified for each of these traits. One accession from Bangladesh, DJ123, which like Kasalath belongs to the *aus* subspecies of rice, combines a favorable phenotype and minor QTL for above three traits (Mori et al. 2016; Wissuwa et al. 2015). DJ123 can therefore be considered a potential candidate donor in breeding programs, with the additional advantage of being an early maturing accession unlike mostly late-maturing landraces from Madagascar.

One of the most successful modern rice varieties in tropical Asia is IR64, released by the International Rice Research Institute (IRRI) in the 1980s. Compared to its predecessors, it combined high yield potential with intermediate tolerance to multiple biotic and abiotic stresses (Mackill and Khush, 2018). Its success in Africa was less auspicious, but it served as recipient variety in several crosses, such as the wide cross with *Oryza glaberrima* donors that gave rise to the lowland NERICA varieties. However, IR64 was never accepted by farmers in Madagascar, and while reasons are not well documented, it is assumed that its semi-dwarf plant type was one deciding factor because farmers prefer longer straw varieties as a source of feed for their cattle and because manual threshing practiced throughout Madagascar is easier with a longer straw plant type. Thus, IR64-derived breeding lines could principally be adapted if donors with taller plant types were used and selection against the short semi-dwarf type was practiced. Such crosses had been made at JIRCAS, Japan, i) between IR64 and medium-tall donor DJ123 and ii) between two BC_2_ IR64-*Pup1* introgression lines of which one also was of medium height.

The objective of this study was therefore to utilize the aforementioned breeding populations in attempts to develop varieties that are adapted to low-input conditions yet are capable of responding to higher soil fertility or moderate rates of fertilizer additions. For this purpose, seed of these two breeding populations was sent to Madagascar, and selections were conducted directly in target environments, which we defined as low soil-fertility farmers’ fields located in villages rather than fertilized fields on breeding stations. Our objective was not to conduct formal participatory variety selection (PVS), however, being located at farmers’ fields and relying on villagers for field operations provided frequent opportunities for farmer feedback during the selection process.

## Materials and methods

### Development of a Pup1 introgression population in the IR64 background

For the introgression of the *Pup1* locus recurrent parent IR64 was crossed with *Pup1*-donor NIL14-4, which contains the *Pup1* region from original donor accession Kasalath in a Nipponbare background (Wissuwa et al. 2005). The resulting F_1_ was backcrossed to IR64 and the BC_1_F_2_ genotyped with *Pup1*-specific markers K20, K29 and K46, where dominant marker K46 is diagnostic of the presence or absence of *OsPSTOL1* and K20 and K29 diagnostic of Kasalath versus IR64 alleles in the *Pup1* region surrounding *OsPSTOL1* (Chin et al. 2011).

The development of *Pup1* breeding lines in the IR64 background is described in detail in Chin et al. (2011). Briefly, a selected BC_1_F_2_ line carrying this introgression was again backcrossed to IR64. BC_2_ progeny from this cross was advanced to the F_3_, phenotyped with K markers (see above), and six homozygous BC_2_F_3_ carriers of Kasalath - *Pup1* were identified. These six lines differed for Kasalath/Nipponbare introgression at loci other than *Pup1* and were selected for further phenotyping: BC_2_F_5_ lines were evaluated for agronomic traits and yield on low-P field plots at IRRI in 2012, and single-plant selections were made. This process was repeated with BC_2_F_n_ (n>6) lines on low-P field plots at Dakawa in Tanzania and on low-P field plots at JIRCAS/Japan in 2014.

Based on these evaluations, two BC_2_F_n_ carriers of *Pup1* were selected: IR64-Pup1-H because of its good yield performance and IR64-Pup1-M because of its yield and larger plant height, which is a trait preferred by many smallholder farmers who utilize straw as animal feed. These two lines were crossed in 2015 to produce a breeding population. Lines were again genotyped for *Pup1* to assure the presence of the introgression and F_3_ seed was sent to Madagascar in 2016 for further selection in P-deficient farmers’ fields.

### Development of a breeding population using donor DJ123 for superior PUE, PAE and P allocation to roots

Crosses between DJ123 and IR64 were made at JIRCAS/Tsukuba in 2015 and progeny advanced to the F_4_ generation by single seed descent. The F_1_ and F_3_ generations were advanced in the greenhouse in small containers whereas the F_2_ and F_4_ – F_6_ generations were grown in a paddy field at the JIRCAS farm during the field season from May to September. F_4_ plants were selected for earliness and acceptable agronomic appearance and in the F_5_ selection was additionally practiced for total panicle weight per plant. From these 50 selected F_6_ lines were shipped to Madagascar in October 2018.

### Selection in the F_3_-F_8_ (IR64-Pup1) and F_5_-F_8_ (DJ123 x IR64) generations

All evaluations of breeding lines were done in farmers’ fields under typical farmer management practices for transplanted lowland rice in Madagascar. Fields were bunded to maintain an even water level, and land preparation (plowing, puddling, leveling) was done by oxen or shovels. No fertilizer was applied to fields, which is common practice at all sites used, nor were other agrochemicals used to control weeds, pests or diseases. Water was supplied by irrigation stemming from small surface creeks, and this was generally sufficient to maintain standing water throughout most of the growing period.

Sowing was done in a seedling nursery located in close proximity to the main field, again without further external inputs. Four-week-old seedlings were uprooted from the nursery and transplanted into the main field as single seedlings with 20 cm spacing within and between rows. Plot sizes varied from a double-row of 10 plants (F_3_) to triple rows of 10 plants (F_5_) to four rows in the F_7_ and 6-row plots in the F_8_. Main selection experiments were conducted in the central highlands during the rainy season from November to May (F_3_, F_5_, F_7_), with sowing typically done in November, transplanting in December and harvesting in April-May. This was followed by a generation of seed multiplication (F_4_, F_6_, F_8_) during the off-season at the coastal site Tsarano/Marovoay (elevation 18 masl, 16°10’52.8”S 46°40’40.0”E), where temperatures and the availability of irrigation from large reservoirs permit year-round rice cultivation. During the off-season, selection was practiced for homogeneity and plant type if progeny was visually segregating. In all trials breeding lines were compared to recurrent parent IR64 and local variety X265 (also known as Malaika), which is the main cultivated variety in the region and appreciated by the farmers. Main selection field trials were conducted in fields located within three villages in the central highlands of Madagascar: Anjiro (elevation 950 masl, 18°54’02.3”S 47°58’11.9”E), Behenjy (elevation 1350 masl, 19°10′48.3″S 47°29′46.3″E), and Ankazomiriotra (elevation 1150 masl, 19°40′07.9″S 46°33′53.9″E).

### Yield trials at farmers’ fields in Madagascar in the 2019-20 and 2020-21 field seasons

Selected breeding lines were evaluated for a potential release as a new variety over 4 cropping seasons between November 2019 and October 2021. Rainy season trials were conducted at farmers’ fields in the central highlands at Anjiro, Behenjy, Ankazomiriotra and Antohobe (elevation 1240 masl, 19°45’60.0”S 46°41’27.6”E) villages with off-season trials being located at two sites in Marovoay. All trials were conducted according to regulations established nationally for Madagascar by the official seed certification agency, the Service Officiel de Contrôle des Semences (SOC; https://soc-semences.mg/homologation-des-varietes/).

Briefly, the process distinguishes DHS trials to evaluate the <u>d</u>istinction, <u>h</u>omogeneity and <u>s</u>tability of a new variety and VATE trials that measure its agronomic value (<u>V</u>aleurs <u>A</u>gronomiques, <u>T</u>echnologiques et <u>E</u>nvironnementales) including tolerance to environmental stresses. DHS evaluations start with a characterization of a candidate variety based on a catalog of plant morphological criteria. These are then confirmed in subsequent seasons during DHS trials, where field plots have a minimum size of 5 x 5 m with single seedlings transplanted at 20 cm distance within and between rows. VATE trials include a NPK (11-22-16) fertilizer treatment with a dose equivalent to 300 kg ha^-1^ and plots were of size 2 x 3 m with 20 cm distance within and between rows. To determine grain yield an area equivalent to 1 m^2^ was harvested. Panicles were separated from culms and air-dried in a greenhouse before weighing. Grain yields are calculated based on a moisture content of 14%.

### Detailed phenotyping experiments

To assess the root angle distribution of crown roots 15 cm diameter metal sieves were buried into a paddy field at the JIRCAS experimental farm in Tsukuba. Japan, so that the top of sieves was about 1 cm below the soil surface and the entire sieve was filled with soil. Twenty-three-day old seedlings of the 5 genotypes were transplanted into these sieves at a depth of about 2 cm. Plants were grown for 5 weeks under irrigated/flooded conditions. A shovel was used to remove the sieves with roots from the field by cutting at about 5 cm from the outside ring so that roots protruding from the sieve were visible. Soil was washed away from the outside of the sieve to be able to count the number of protruding crown roots in four classes depending on their angle: i) shallow surface roots above the rim of the sieve (< 1°); ii) root protruding from the top 3 cm of the sieve (1-25°); iii) roots below 3 cm but above the flat sieve bottom (25-60°); iv) roots protruding from the sieve bottom (>60°) (see Figure 4b).

For the evaluation of genotypic differences in PUE plants were grown hydroponically in a greenhouse with natural light at temperatures set to 30°C during the day and 24°C at night. Seeds were germinated in petri dishes and after germination were transferred to a mesh floating above a solution containing 0.1 mM calcium (CaCl) and 18 μM iron (FeEDTA). Two-week-old seedlings were then transferred into experimental units: 1-L black plastic bottles containing half strength Yoshida solution (Yoshida et al. 1976) in which the P concentration was reduced to 3 µM (KH_2_PO_4_). The nutrient solution was replaced in weekly intervals and for the 4^th^ and 5^th^ week the Yoshida solution was increased to full-strength except for the constant 3 µM P. Thus, plants received 4 additions of 3 µmol P for a total supply of 371.6 µg P per plant. Plants were harvested when they were 6 weeks old, root and shoot was separated, oven-dried for 4 days at 60°C, and weighed. The P concentration in roots and shoots was determined after digestion in a mixture of HNO_3_ and HClO_4_ (3:1), followed by P concentration measurements using the molybdenum blue method (Murphy and Riley, 1962).

A second nutrient solution experiment was conducted in a greenhouse at the University of Bonn under natural light and temperatures set to 30°C during the day and 24°C at night. The pre-treatment followed the protocol described above. Eight days after germination 5 seedlings per genotype were transferred into 15-L plastic boxes that had 20 holes drilled into the lid to fasten seedlings with foam strips. The boxes contained initially ¼ strength Yoshida solution with either high (50 µM P) or low (1 µM P) P supply. On day 5 Yoshida stock solution (equivalent to 0.2X) was added together with an addition 1 µM or 50 µM P. On day 10 the solution was replaced with 0.5X Yoshida solution and the P concentration in the low-P treatment increased to 2 µM P. A final addition of 0.25X Yoshida and 2 or 50 µM P was given on day 14 and plants were harvested on day 18, crown roots counted, roots and shoots separated and oven-dried for 4 days at 60°C.

### Whole-genome resequencing and data analysis

Genomic DNA from selected breeding lines, Kasalath, IR64, and DJ123 was extracted using the standard phenol/chloroform method using a leaf of a single plant. A sequencing library was created with the TruSeq DNA PCR Free kit (Illumina) and sequenced on a NovaSeq6000 (Illumina), generating 150 bp of paired-end reads. The raw reads were analyzed and aligned to the IR64 reference genome (GCA_009914875.1) as reported previously (Pariasca-Tanaka et al. 2025). The resulting vcf file was thinned to provide genotype information for every 20 kb using TASSEL5 software (Bradbury et al. 2007).

## Results

### Variety development of FyVary32 and FyVary85

Details of the selection process are summarized in Figure 1. F_3_ progeny lines derived from a cross of two BC_2_ IR64-Pup1 introgression lines (IR64-*Pup1*-M x IR64-*Pup1*-H) were grown in an unfertilized farmer’s field at Anjiro village. From the 38 segregating F_3_ families 80 individual plants were selected based on earliness (maturing earlier than IR64), vigor, and panicle weight per plant (May 2016). Since the amount of grain produced under zero-input conditions was limited, seed of selected plants was increased for one generation during the off season on a less P deficient irrigated site in coastal Marovoay. Evaluations of the 80 F_5_ lines were conducted at three field sites in Anjiro, Behenjy, and Ankazomiriotra villages and 21 plants were selected (May 2018). One of these was de-selected during seed multiplication in the off-season, and the remaining 20 were evaluated again in the F_7_ at the same three villages (Nov 2018 – May 2019). From these trials, seven IR64-*Pup1* lines were selected in May 2019 for seed multiplication (F_8_) and variety release testing in the F_9_ generation.

**Figure 1.**
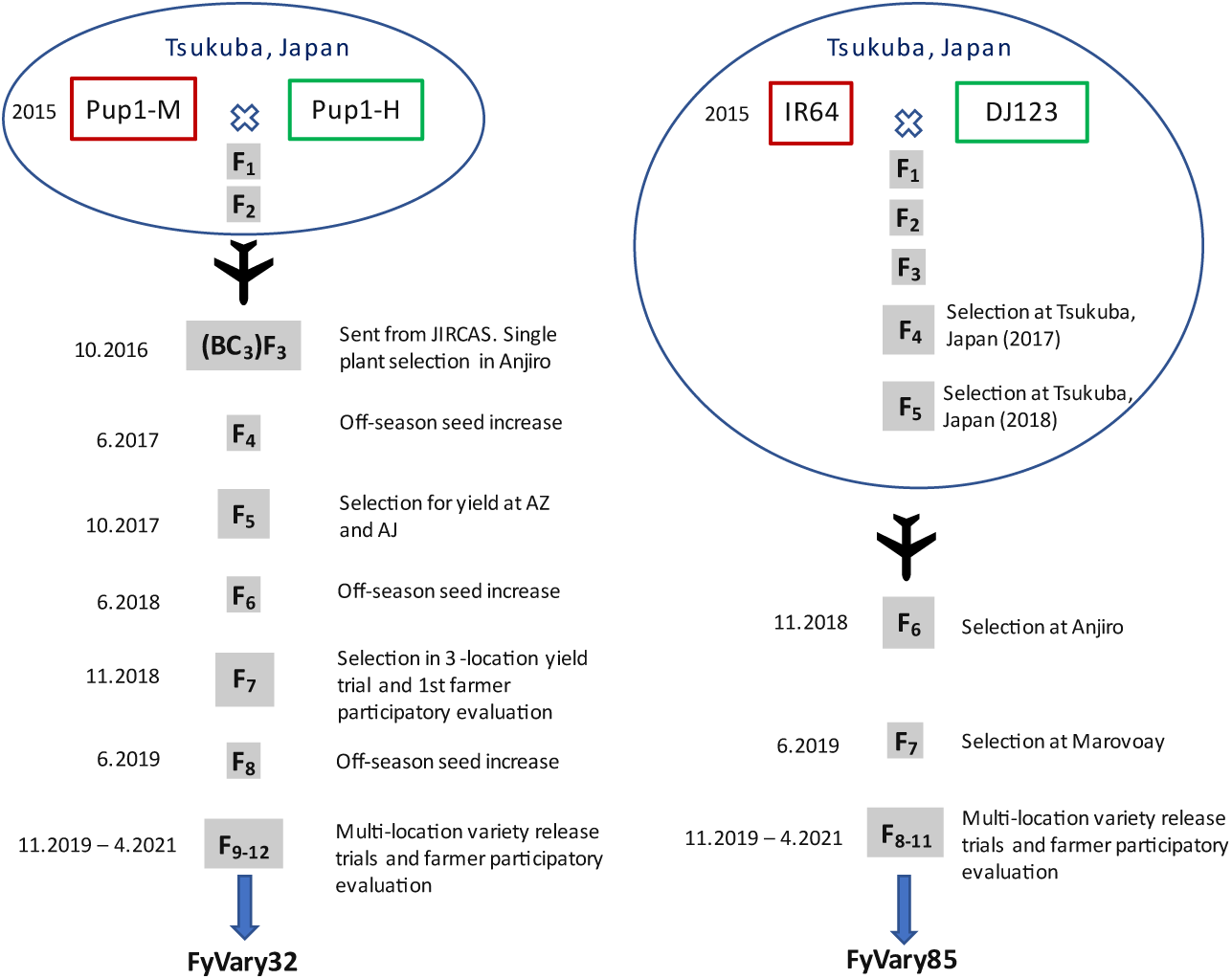
Development scheme for varieties FyVary32 and FyVary85. Crosses and early stages were done at JIRCAS, Japan. Breeding lines were then sent to Madagascar for selection in farmers’ fields under low input conditions. The variety release evaluation was conducted jointly for all breeding lines from November 2019 until April 2021.

A 2^nd^ breeding population was developed from the cross of IR64 and DJ123, a donor for high PUE and RE, at JIRCAS, Japan, from 2015 onwards. In the F_4_ (2017) and F_5_ (2018) generations selection for biomass and total panicle weight was practiced at the JIRCAS experimental farm on a low-input field. 50 selected F_6_ lines were sent from JIRCAS, Japan to Madagascar in October 2018. Plants were grown in an unfertilized farmers’ field at Anjiro village (2018-19 season) and 11 individual plants were selected based on superior biomass and grain weight per plant. After further evaluation during the off-season in 2019, 3 selected F_8_ lines were included in variety release testing starting in the 2019-20 main season of which only line #85 was advanced to the 2^nd^ year.

All selection experiments were conducted at farmers’ fields with hired labor from the surrounding village. Farmer feedback on the breeding material was frequently obtained and considered during the selection process. In addition to these informal participatory selection inputs, formal PVS evaluations were included during the variety release procedure.

### Yield trials at farmers’ fields in Madagascar in the 2019-20 and 2020-21 field seasons

Official variety evaluation trials according to Malagasy government regulations and supervised by the Service Officiel de Contrôle des Semences (SOC). The process distinguishes DHS trials to appraise the <u>d</u>istinction, <u>h</u>omogeneity and <u>s</u>tability of a new variety and VATE trials that evaluate its agronomic value (<u>V</u>aleurs <u>A</u>gronomiques, <u>T</u>echnologiques et <u>E</u>nvironnementales), including tolerance to environmental stresses. Furthermore, hedonic tests to gauge consumer acceptance were included with at least 100 consumers in each of the four test villages. In all trials, breeding lines were compared to the parental variety IR64 and to local check X265, which is the recommended variety for the central highlands of Madagascar. These trials started in the main season 2019-20 and continued for 2 main and 2 off-seasons (Figure 1) until a final evaluation in October 2021 and the eventual release as FyVary32 (from IR64-*Pup1* line #32) and FyVary85 (from DJ123 x IR64 line #85) in November 2021. Initially seven IR64-*Pup1* lines and three DJ123 x IR64 lines had been included but four IR64-*Pup1* lines were omitted after the first year and two after the 2^nd^ year (lines #45 and #52). This 2^nd^ omission was due to lines #45 and #52 being not distinct enough from #32.

Over the 18 trials conducted the highest average grain yield was achieved by FyVary85 (3.97 t ha^-1^), followed by FyVary32 with 3.67 t ha^-1^ (Table 1). This represented an improvement over X265 of 19.6% and 11.7%, respectively, and of 22.2% and 12.9% over parental variety IR64. The superiority of FyVary85 increased further at less productive sites with average grain yields below 3 t ha^-1^. Here FyVary85 surpassed X265 by 25.7% and parent IR64 by more than 30% (Table 1). At sites exceeding 3.8 ha^-1^, which included the same low-fertility sites after addition of NPK fertilizer (n=5) as well as sites with higher native soil fertility (n=8), FyVary32 and 85 exceeded X265 significantly (12.4 – 14.9%), with a slightly more pronounced advantage over parent IR64 (14.7 – 17.2%).

**Table 1.**
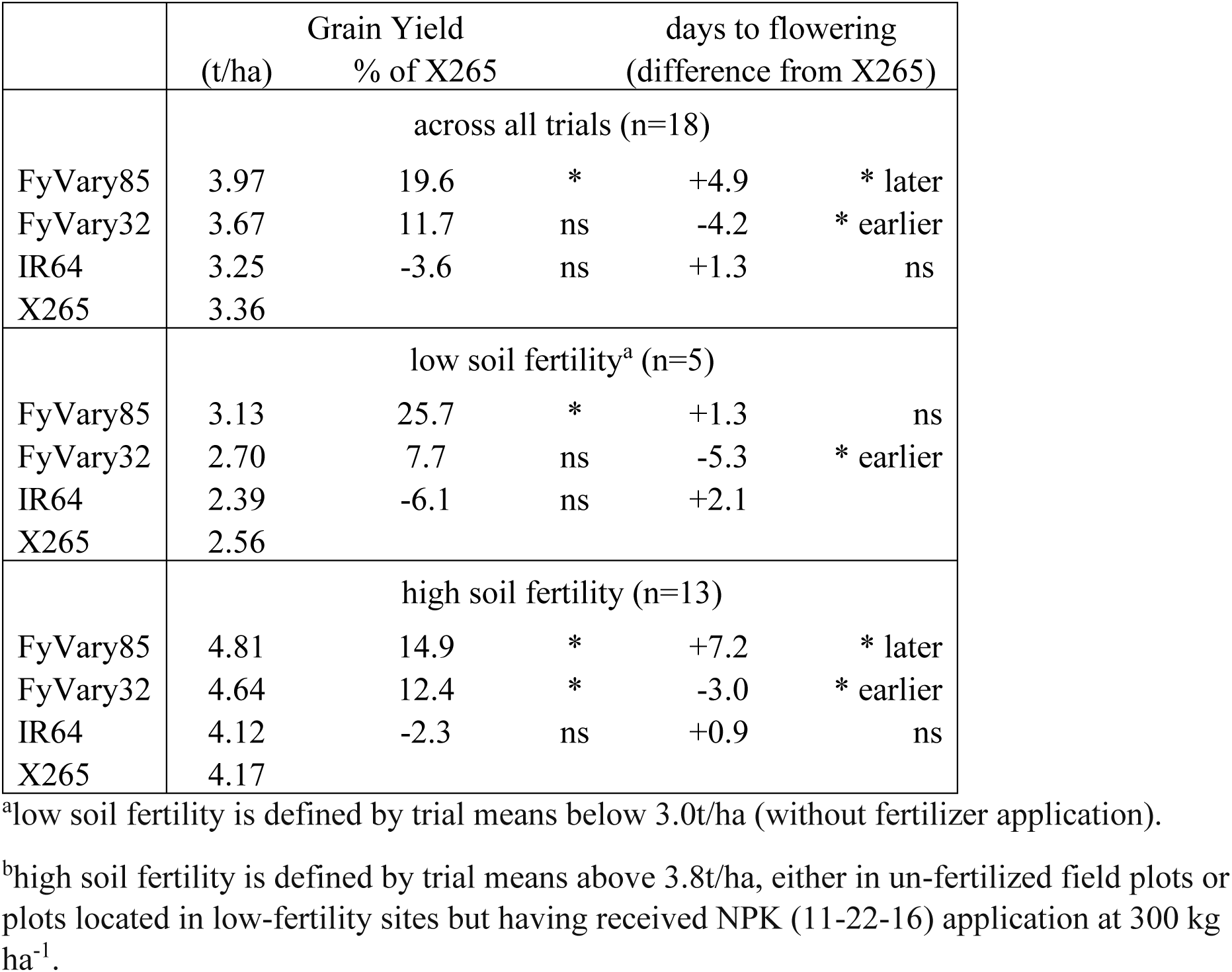
Grain yield and days to flowering in newly released varieties FyVary32 and 85 compared to local check X265 and parental variety IR64. Trials were conducted over 4 seasons from November 2020 to October 2021 at five villages in Madagascar.

|  | Grain Yield |  | days to flowering |  |  |
| --- | --- | --- | --- | --- | --- |
|  | (t/ha) | % of X265 |  | (difference from X265) |  |
|  | across all trials (n=18) |  |  |  |  |
| FyVary85 | 3.97 | 19.6 | * | +4.9 | * later |
| FyVary32 | 3.67 | 11.7 | ns | -4.2 | * earlier |
| IR64 | 3.25 | -3.6 | ns | +1.3 | ns |
| X265 | 3.36 |  |  |  |  |
|  | low soil fertility<sup>a</sup> (n=5) |  |  |  |  |
| FyVary85 | 3.13 | 25.7 | * | +1.3 | ns |
| FyVary32 | 2.70 | 7.7 | ns | -5.3 | * earlier |
| IR64 | 2.39 | -6.1 | ns | +2.1 |  |
| X265 | 2.56 |  |  |  |  |
|  | high soil fertility (n=13) |  |  |  |  |
| FyVary85 | 4.81 | 14.9 | * | +7.2 | * later |
| FyVary32 | 4.64 | 12.4 | * | -3.0 | * earlier |
| IR64 | 4.12 | -2.3 | ns | +0.9 | ns |
| X265 | 4.17 |  |  |  |  |

<sup>a</sup>low soil fertility is defined by trial means below 3.0t/ha (without fertilizer application).
<sup>b</sup>high soil fertility is defined by trial means above 3.8t/ha, either in un-fertilized field plots or plots located in low-fertility sites but having received NPK (11-22-16) application at 300 kg ha<sup>-1</sup>.

The number of days between sowing and flowering varied by more than 30 days between sites (data not shown), with average temperatures, degree of soil fertility and fertilizer applications likely being the main determining factors. To be able to summarize genotype effects for days to flowering despite such large site-specific variation, the difference in days compared to X265 at each site was used instead of the total period. Across all sites FyVary32 flowered 4-5 days earlier than X265 and IR64 and that increased slightly at sites with a lower yield level (Table 1). FyVary85 was generally flowering later than X265 but this was much less pronounced on sites with low yield potential compared to higher-yielding sites.

The responsiveness of FyVary varieties to increasing soil fertility was examined by plotting line mean grain yields at a site against the site mean grain yield (Figure 2). Site mean yield ranged from as low as 2.12 t ha^-1^ at Anjiro (2021) to as high as 5.58 t ha^-1^ at a second Anjiro site that received NPK fertilizer. At each site one of the FyVary varieties was the highest yielder (10 times FyVary85 and 8 times FyVary32). Comparison of the fitted regression lines indicated that differences in responsiveness were very minor and that FyVary85 had a yield advantage over X265 of 0.638 t ha^-1^ across the entire range. FyVary32 and IR64 were slightly more responsive with coefficients of 1.038 and 1.051, respectively, and with this similar responsiveness FyVary32 had about 0.47 t ha^-1^ higher grain yield compared to IR64.

**Figure 2.**
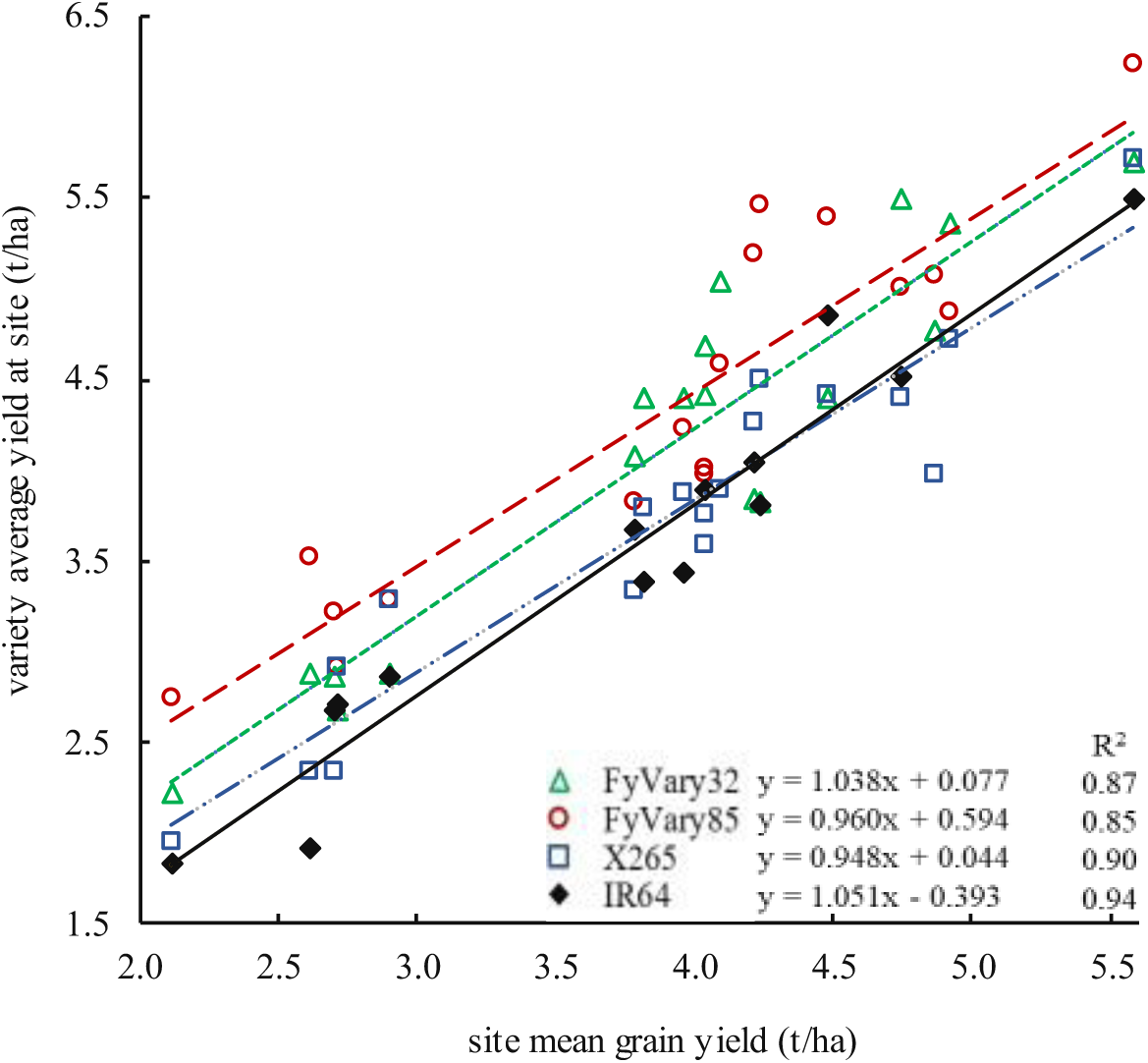
Grain yield of newly released varieties FyVary32 and 85 compared to parental variety IR64 and local check X265 during variety release trials in 2019-2021. Average grain yields of varieties at a site are plotted against the site mean yield on the x-axis. Linear regressions of variety mean yields versus site mean yields are presented including the corresponding R^2^.

FyVary32 was significantly and up to 20 cm taller compared to IR64 (Figure 3) and thus can no longer be considered of semi-dwarf type whereas FyVary85 is intermediate and appears more sensitive to changes in fertility. FyVary varieties are contrasting for panicle length with FyVary32 having a long and slender panicle while FyVary85 is characterized by a short and dense panicle (Figure 3C). Both varieties produce more productive tillers, however, the differences was only significant compared to X265 but not to IR64. For thousand grain weight genotypic differences were not significant.

**Figure 3.**
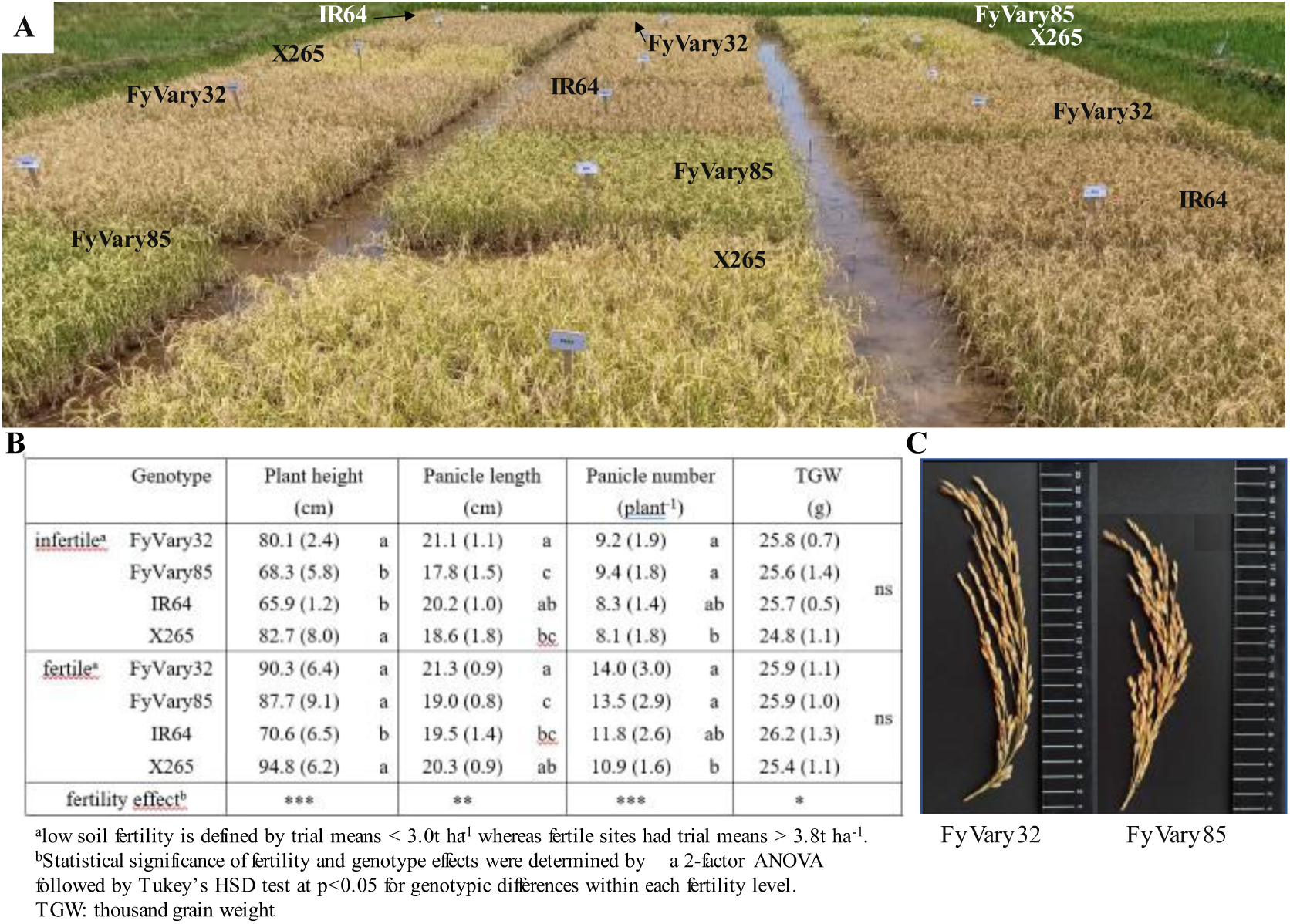
(A) FyVary32 and FyVary85 in the final variety release trial managed by FOFIFA in Marovoay village, together with check varieties X265 and IR64 and two sister lines (#45 and #52) that were not released. (B) Plant height, panicle length and number, and thousand grain weight (TGW). Data shown are means (standard deviation) over trials conducted between November 2020 to October 2021. (C) Panicles of FyVary32 and FyVary85.

**Figure 4.**
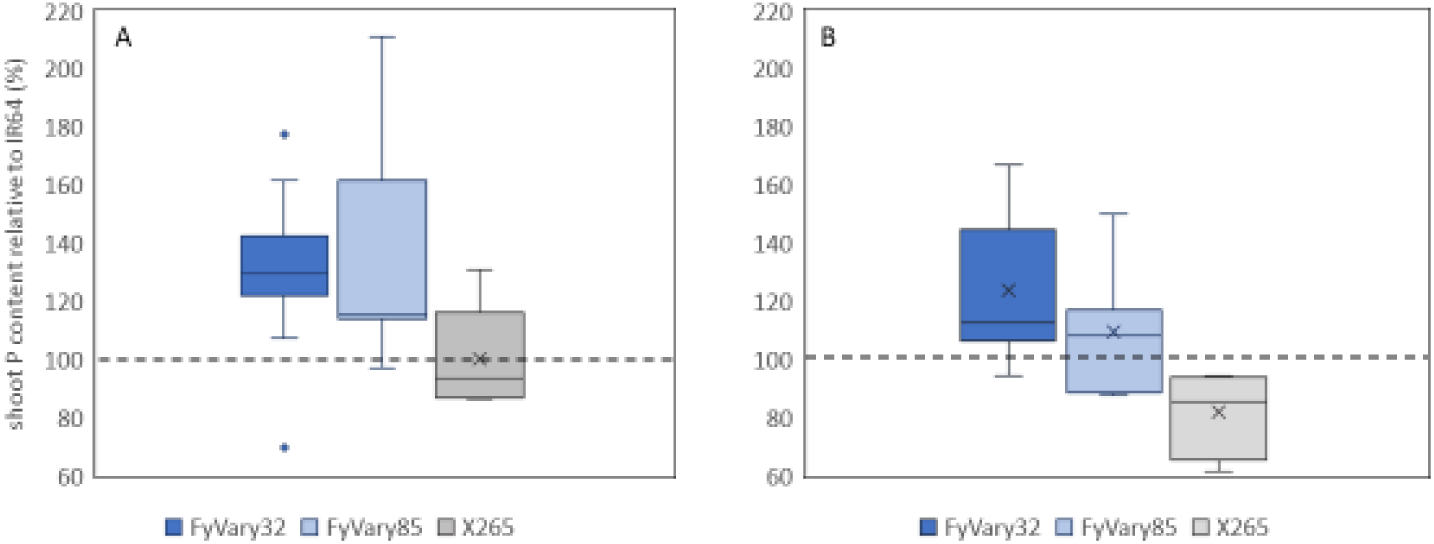
Shoot P content of FyVary varieties compared to IR64 and X265 at 5 P deficient sites (A) and 2 sites with intermediate P level (B). Data is expressed relative to the P content of IR64 at each site (in %).

Conducting a hedonic test is a prerequisite for a successful variety release. As part of participatory varietal selection (PVS), this test is essential because farmers’ appreciation of taste strongly influences their acceptance of a new variety. It was conducted at 4 villages with >100 participants each. Rice was prepared locally by villagers and their impression on appearance, taste and texture recorded by staff of the National Center for Applied Research on Rural Development (FOFIFA). Slight differences in preference existed between villages, with FyVary32 being preferred in Anjiro compared to FyVary85 in Marovoay (suppl. Figure S1). However overall scores only ranged between 3.61 – 3.93 (on a scale of 1 – 5; 5 being best) and rankings of both varieties were not significantly different from local variety X265.

### Effects of selection on P uptake, efficiency and root growth

P uptake was evaluated at 5 sites during the seedling stage about 4-5 weeks after transplanting. To compare data across sites with different soils and growth potential, FyVary varieties were compared relative to IR64 at each site (IR64 = 100). At sites categorized as P deficient average shoot P concentrations ranged from 1.04 – 1.26 mg g^-1^ compared to 1.51 - 1.62 mg g^-1^ at sites with intermediate P availability. Highest P uptake relative to IR64 was seen in FyVary85 with 137.6% with FyVary32 at 130.5% (Figure 4). In the intermediate P sites average biomass had doubled and P uptake increased 2.5-fold relative to P deficient sites (data not shown) and under these more favorable conditions the superiority in P uptake over IR64 decreased to 23.9% for FyVary32 and 9.3% for FyVary85.

### P efficiency traits in FyVary varieties compared to parent IR64 and X265

Based on the characteristics of donors used in crosses with IR64 it is expected that resulting FyVary varieties may possess a combination of improved P-efficiency traits such as increased crown root number conferred by the *Pup1* locus, a generally larger root system or higher internal PUE (from donor DJ123).

Crown root number was evaluated in two of the 7 sites used to determine P uptake and FyVary85 and FyVary32 had significantly more crown roots per plant (67.7 and 61.2, respectively) compared to IR64 with 50.3 crown roots per plant (Figure 5A). To further characterize the root system of FyVary varieties compared to their parents and X265, the root basket method was applied in an irrigated high-fertility field at the JIRCAS experimental farm (Figure 5B). With 390 total crown roots FyVary85 had significantly more roots compared to remaining genotypes which were not different and ranged from 262 in IR64 to 279 in X265 (Figure 5C). DJ123, the parent of FyVary85, differed from all other genotypes in its distribution of crown root angles as more that 70% of roots had an angle steeper than 25° (Figure 5C). Parent IR64 and X265 on the other hand were characterized by a very shallow root system with about 70% of roots having an angle < 25°. FyVary varieties were intermediate with a significantly higher number of roots between 25° to 60° compared to IR64. FyVary85 had the most balanced root system and due to its very high total root number, it was among the group with highest root number in the surface, shallow and intermediate classes. Only DJ123 had a significantly higher number of deep roots >60° than FyVary85.

**Figure 5.**
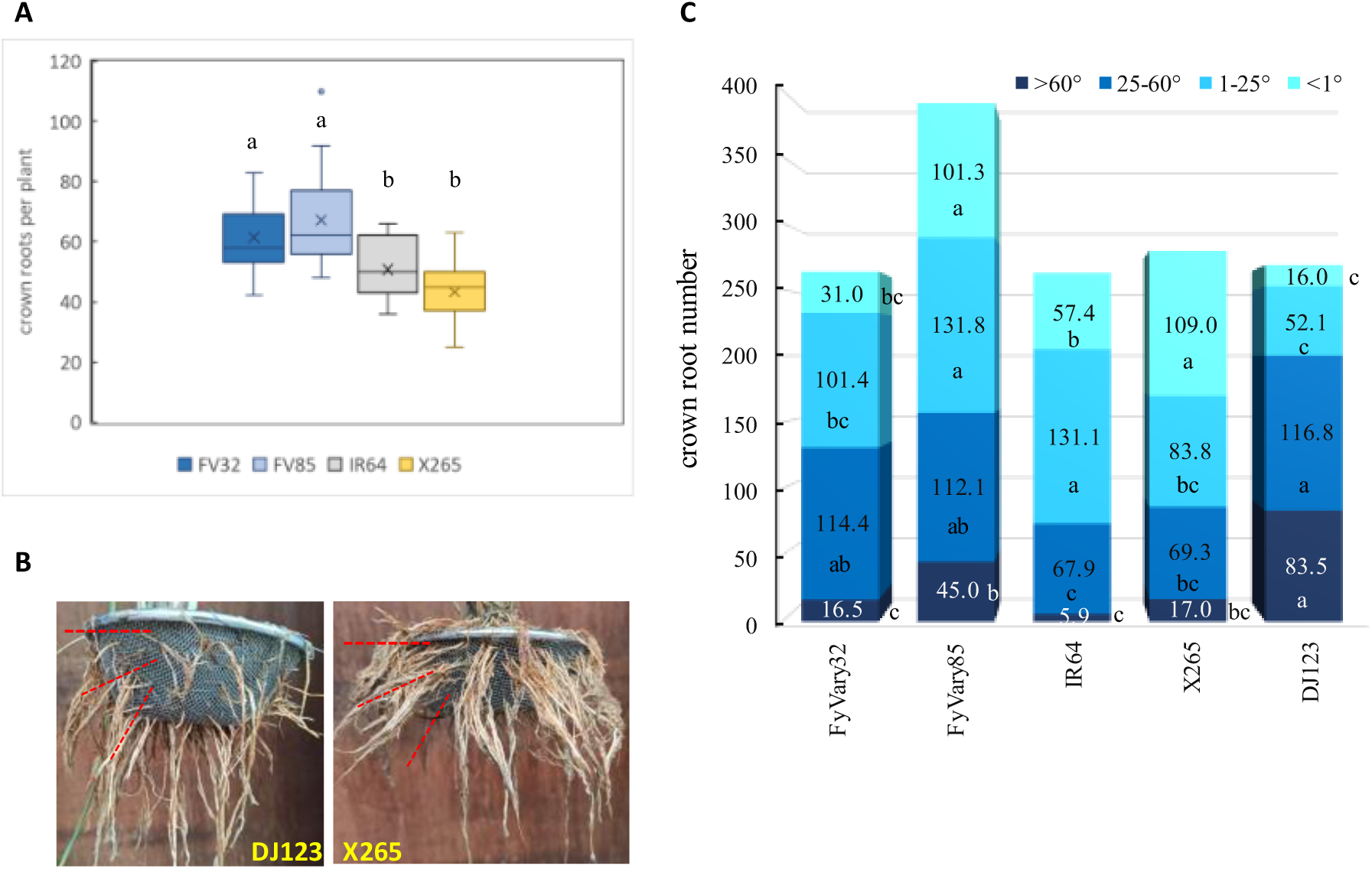
(A) Crown root counts per plant from plants dug out at tow field sites located in Marovoay town, Madagascar. Different letters indicate significant differences according to Tu ey’s SD test at p 0.05. (B) The basket method used to determine root angle distributions where roots protruding from the basket at different angles (< 1°, 1-25°, 25-60°, and >60°; indicated by dashed lines) were determined after excavating the baskets from a paddy field located at the JIRCAS experimental farm. (C) The distribution of roots in 4 classes were determined where class <1° corresponds to roots growing at the soil surface above the basket.

The internal P utilization efficiency (PUE) was examined in a nutrient solution experiment where each plant was grown in its own 1.1-L container to which a total of 372 µg P had been added. Total P content did not differ between varieties (data not shown), indicating that the design purpose of testing varieties at the same P-deficiency stress level (same plant P content) was achieved. FyVary85 was using P most efficiently, producing 2.25 and 2.53 g biomass per mg P in shoot and root tissue, respectively (Table 2). The difference to parent IR64 was significant in shoot (+20%) but not root tissue (+7%) whereas FyVary32 did not differ from IR64.

**Table 2.** Internal P utilization efficiency (PUE) in shoot and root tissue with respective biomass in a nutrient solution experiment where each plant was supplied with 372 µg P.

|  | PUE (g biomass mg<sup>-1</sup> P) |  |  |  | Biomass (g plant<sup>-1</sup>) |  |  |  |
| --- | --- | --- | --- | --- | --- | --- | --- | --- |
|  | shoot |  | root |  | shoot |  | root |  |
| FyVary32 | 1.92 (0.07) | c | 2.42 (0.12) | ab | 0.564 (0.039) | b | 0.354 (0.019) | ab |
| FyVary85 | 2.25 (0.05) | a | 2.53 (0.05) | a | 0.633 (0.021) | a | 0.380 (0.004) | a |
| IR64 | 1.87 (0.06) | c | 2.36 (0.10) | ab | 0.518 (0.020) | b | 0.322 (0.018) | b |
| X265 | 2.10 (0.08) | b | 2.29 (0.09) | b | 0.581 (0.042) | b | 0.322 (0.018) | b |

Genotype means followed by different letters are significantly different at p<0.05 according to Tuckey's HSD test.

In a second nutrient solution experiment crown root number and biomass was evaluated in 24-day-old plants grown under high and low P supply. The low P supply reduced total dry weight (TDW) by 26.4% in IR64 with less than 10% reductions in FyVary 32 and 85 (Table 3). Under high P supply biomass accumulation did not differ significantly between genotypes but under P deficient condition FyVary85 maintained the highest total biomass, followed by FyVary32 and IR64. With high P supply IR64 produced the largest number of crown roots but this reversed under P deficiency with FyVary32 being the genotype with significantly more crown roots (Table 4). Remarkably crown root number increased slightly under P deficiency in FyVary32 whereas crown root number was reduced by 20% in IR64.

**Table 3.**
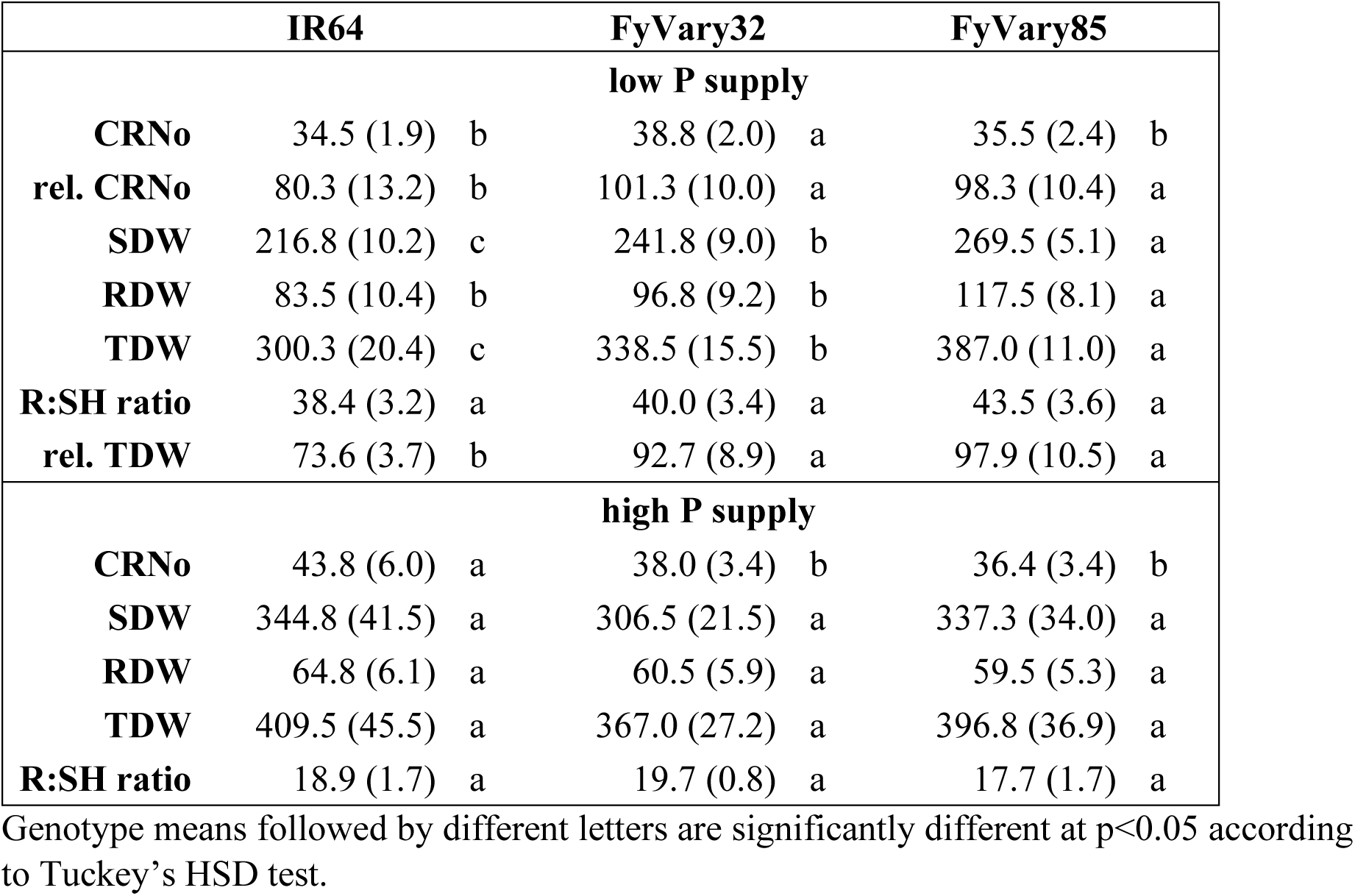
Crown root number (CRNo) and biomass traits in IR64 compared to FyVary varieties. Following a 8-day pre-treatment period without P supply plants were grown for 18 days in nutrient solution with low (2 μM) or high (50 μM) P supply.

### Graphical genotype of FyVary32 and Pup1 genotype of FyVary85

To identify donor introgressions other than the Pup1 locus the graphical genotype of FyVary32 was obtained by aligning short read sequences to the IR64 reference sequence (GCA_009914875.1). This confirmed that FyVary32 contained a large donor introgression at the *Pup1* locus between 9.48 – 16.59 Mb on chromosome 12 (suppl. Figure S2A). Additional introgressions supported by multiple SNPs were detected on chromosomes 1, 2, 5, 6, and 11. The introgression on chromosome 1 between 38.2-40 Mb contains the Nipponbare allele at the SD1 locus and is likely responsible for the 14 – 20 cm larger plant height (see Table 2).

Using *Pup1*-specific markers we furthermore showed that FyVary85 inherited the *Pup1* region from parent IR64, which meant that the large indel containing *PSTOL1* was absent in FyVary85 (suppl. Figure S2B).

## Discussion

The need to boost rice production is SSA has been highlighted repeatedly and several internationally supported initiatives such as the Coalition for African Rice Development (CARD) or the Alliance for a Green Revolution in Africa (AGRA) have been initiated over the past decades. However, attempts to replicate the Asian green revolution in SSA were of very limited success and increases in overall rice production have been largely due to an expansion of the land area cropped with rice while yields remained stagnant (Yuan et al. 2024) or increased mainly in the relatively few irrigation schemes that allow for more targeted interventions (Otsuka et al 2024). Outside such islands of higher productivity yield gaps remain large, often exceeding 80% of attainable yields (Yuan et al. 2024).

Frequently, one argument put forward for the need to increase rice productivity is the widening gap between country level rice demand, which is increasing rapidly across SSA including Madagascar, and country level rice production (Ashtakala et al., 2025; Dröge et al., 2022; Yuan et al., 2024). Negative effects of climate change will exacerbate this gap further, thereby increasing food insecurity in Madagascar (Dröge et al., 2022). The expansion of irrigation facilities and adoption of the package of inorganic fertilizers, improved agronomic practices and modern varieties (MVs) in more favorable environments is seen as a possible solution to prevent a widening gap between demand and local supply (Otsuka et al 2024). In addition to the country-wide needs to boost production, which may target certain regions of strategic relevance, efforts to boost rice production should also address the goal of improvements in household food security, which is of particular relevance for smallholder farmers (Tojo-Mandaharisoa et al, 2022) that frequently do not produce sufficient rice to feed the household until the coming harvest (Minten and Barrett, 2008). Many of these farmers are located in less favorable ecologies such as upland areas or small inland valleys, both characterized by weathered, infertile soils and reliance on rainfall rather than irrigation (Andriamananjara et al., 2018). In this context, a more differentiated approach in relation to green revolution technologies may be required, and we argue that this should also include breeding targets.

### A case for a dual approach to intensification that includes alternative breeding objectives

In the selection of varieties during the GR emphasis has mainly been on achieving high yield potential (Mackill and Khush, 2018), and in the years since this focus has been maintained while also targeting tolerance to biotic and abiotic stresses. Selections and breeding line evaluations have typically been conducted on research or breeding stations under very favorable conditions that include the application of inorganic fertilizers at recommended rates to assure the yield potential can be assessed. Subsequent testing in multi-environment trials may have included less favorable sites, but the omission of inorganic fertilizers to test for adaptation to low-input conditions has generally not been an objective. Furthermore, other constraints experienced by farmers, such as limited financial resources to provide optimal inputs and extreme agroclimatic variability, are often disregarded (Weltzien & Christinck, 2017). Thus, released varieties from international centers like IRRI and AfricaRice, and from a few strong national rice breeding programs present in SSA, tend to possess high yield potential, high responsiveness to fertilizer inputs and tolerance to some biotic and abiotic stresses. In addition, preferences are often given to populations that perform well under a wide range of conditions, especially with regards to breeding economics (Weltzien & Christinck, 2017). Their adoption by rice farmers across SSA is, however, low, and often there are insignificant yield increases under farmers’ production conditions, highlighting the unsuitability of these varieties under farmers’ conditions (Mujawamariya et al., 2022; Weltzien & Christinck, 2017) with more likely adoption by farmers in favorable conditions and with financial means and easy access to fertilizers. In contrast, smallholder farmers are more likely retain traditional varieties and use their own produced seeds (Weltzien & Christinck, 2017).

This clearly indicates that modern rice breeding has not satisfied the needs of many smallholder rice farmers, nor has it accounted for the complexity of production goals and the importance of farming for certain people’s livelihood strategies (Weltzien & Christinck, 2017). Farmers may value different attributes than researchers since a crop can fulfil multiple functions in the farming system (Weltzien & Christinck, 2017). In Madagascar, for example, farmers practice mixed crop-livestock farming and therefore value taller varieties that produce sufficient straw for animal feed (Fanjaniaina et al., 2021).

This study has addressed the issue of how to also bring breeding progress to these smallholder farmers. In our approach, we have deviated from typical rice breeding in the choice of the selection environment, which we have shifted from station to farmers’ fields, adopting farmer management practices with regard to land preparation, water management and fertilizer inputs. None of the fields used for selection received inorganic fertilizers, while some fields had received organic fertilizers (composted cattle manure) in preceding seasons. We deviated from farmers’ practice in planting density. Instead of transplanting around 5 seedlings per hill at 12-15 cm spacing between hills within and between rows as is common on farms, we transplanted single seedlings and maintained the 20 cm spacing within and between rows typically used in on-station breeding nurseries in order to facilitate selection of homogenous lines. During the selection process (i.e. the F_4_ – F_8_ generations), no formal participatory variety selection (PVS) was practiced; however, since local farmers were hired for all field operations, their opinions or comments were frequently invited and considered in selections.

### Breeding of FyVary32 and FyVary85 – modern varieties adapted to low soil fertility

This low-tech farm-centered approach employed during selection contrasts with the much more research-driven, formal approach in our donor selection and population development. The development of FyVary32 involved MAS for the *Pup1* locus mapped in 2002 (Wissuwa et al. 2002) and the underlying *OsPSTOL1* gene subsequently identified through positional cloning (Gamuyao et al. 2012). The MAS practiced here differed from other instances like MAS for the *sub1* locus (Neeraja et al. 2007) in as much as the target was not to fully reconstitute IR64 through repeated backcrossing, because IR64 itself has not been adopted widely by farmers in Madagascar or elsewhere in SSA. In the case of Madagascar, one assumed reason is its semi-dwarf plant type that does not produce sufficient straw biomass to satisfy farmers’ needs as animal feed. Our approach, therefore, was to fix *Pup1* in IR64-derived breeding lines in the BC_2_ generation and to further select for more suitable plant types based on the remnant variation present in BC_2_ lines. Evaluations of these BC_2_ lines then identified the two parents used in the final cross from which FyVary32 was selected: IR64-Pup1-H and IR64-Pup1-M, both with good yield performance and the latter with 15-20 cm larger plant height compared to IR64, which was retained in FyVary32 with a 14-17 cm larger plant height compared to IR64.

P-efficiency related traits other than *Pup1* were targeted in the development of FyVary85. Donor accession DJ123 was identified through phenotyping a GWAS panel of IRRI gene-bank accession for internal P utilization efficiency (Wissuwa et al. 2015) and external P acquisition efficiency (Mori et al. 2016). More detailed investigations confirmed high PUE and PAE in DJ123 as it produced more biomass per amount of P taken up (Wissuwa et al. 2015) and acquired more P per unit root size from a P-fixing soil compared to a modern variety (Wissuwa et al. 2020).

In FyVary 32 crown root number in field trials increased compared to recurrent parent IR64 as expected after the introgression of the *Pup1* locus. Experiments conducted in nutrient solution furthermore indicated that the difference in crown root number was not constitutive but specific to P deficient conditions where it was significantly reduced in IR64 but not in FyVary32. FyVary85 does not contain the *Pup1* locus but nevertheless produced the highest crown root number in field trials, which must have been conferred by loci other than *Pup1* and *qCRN9* previously identified from donor DJ123 (Dinh et al 2023) is one possible candidate. DJ123 is furthermore a donor of high internal PUE, and this trait was inherited by FyVary85, producing 20% more shoot biomass per unit shoot P content compared to IR64. The presence of multiple P efficiency mechanisms in FyVary85 is the likely explanation for its increasing superiority over IR64 in P uptake and grain yield under decreasing soil fertility.

Both varieties were selected in farmers’ fields without added mineral fertilizers, representing typical smallholder farmers’ conditions with expected yield levels ranging from around 2.5 – 3.5 t ha^-1^. During variety release testing, the range of site yields expanded to 2.2 – 5.5 t ha^-1^, with higher yields being achieved in a few higher-fertility sites and where NPK fertilizers were applied to fulfil national variety release procedures. Despite selection not being conducted under such conditions for both varieties, they maintained higher grain yields compared to IR64 over the entire range. For FyVary32, the response to better soil fertility was highly similar to IR64, maintaining a yield advantage of about 0.47 t ha^-1^ for the entire range. FyVary85 also surpassed IR64 for the entire range, but its superiority was highest (0.7 – 0.8 t ha^-1^) in the lowest yielding environments. We did not consider evaluations in highly productive environments because these are very rare exceptions in the country and therefore not practically relevant. Our example shows that selection environments at the lower end of expected soil fertility did not compromise yield potential at the higher end of the relevant yield range, and this is likely due to the choice of IR64 as the recipient parent, which produced varieties largely resembling a modern rather than a traditional variety.

The choice of suitable selection environments has been discussed repeatedly, with selection in uniform high fertility environments typically found on breeding stations having the advantage of achieving greater heritability (Atlin et al 2001; Dawson et al 2008; Castro-Pacheco et al 2024). High on-station heritabilities are, however, not practically relevant in the presence of cross-over interactions for grain yield and low genetic correlations between high fertility and moderate to low fertility environments, and such low genetic correlations have been repeatedly detected (Atlin et al 2001), including for rice in Madagascar (Castro-Pacheco et al 2024). While our study was not designed to examine cross-over interactions between high and moderate to low fertility environments, we can conclude that selection in the low to medium fertility environments did not produce crossover interaction in our varieties up to what can be considered high-fertility environments in the context of Madagascar (i.e. almost twice the national average grain yield).

### The potential impact of FyVary varieties

FyVary32 and 85 were released in November 2021, and seed production commenced in the 2021-22 season in collaboration with FOFIFA. Through efforts of the JICA (Japan International Cooperation Agency) funded PAPRIZ project and in collaboration with FOFIFA, foundation seed was made available to more than 50 local seed producers in 13 of the 24 regions of Madagascar, which had produced 31t and 33t of certified seed of FyVary32 and 85, respectively, and feedback from producers indicated the entire stock was sold to farmers at the start of the 2023-24 season (PAPRIZ, pers comm.). Unfortunately, no further official data on seed quantities produced and area grown is available. One limitation in Madagascar and likely in much of SSA is that the formal seed sector is very underdeveloped. The relatively high costs for official seed inspections and certification likely mean that much of the produced seed is sold in the informal seed market, spreading slowly from intervention points as those previously managed through internationally funded projects (i.e. the PAPRIZ project). In recognition of the possible impact initial outside funding may have on the spread of new technologies such as FyVary seeds, a new project was initiated between JICA and the Ministry of Agriculture in Madagascar to produce and distribute at least 100t of certified FyVary seed in the coming years (JICA 2024). This will be bundled with efforts to spread fertilizer micro-dosing options as an entry point into sustainable intensification of smallholder rice production in Madagascar (Tsujimoto 2025).

## Conclusions

We argued for a dual approach in rice breeding that not only targets the development of high-yielding varieties adapted to favorable environments but would also allow smallholder farmers in more marginal environments to benefit from advances in crop breeding. Our example with FyVary varieties indicates that pre-breeding research conducted far from target environments, focusing on identifying tolerance alleles and on targeted exploitation of gene-bank resources, can be harnessed for the benefit of smallholder farmers when selection and variety development stages are moved to a target environment that represents farmers’ conditions and constraints. The constraint we addressed is low soil fertility with P deficiency caused by high P-fixation in highly weathered tropical soils, being the main limitation. In the literature, the approach of selecting in a target environment or decentralized breeding is often contrasted with breeding for broad adaptation in centralized formal breeding approaches. However, as pointed out by

Dawson et al (2008), this presumed broad adaptation can be misleading if it merely refers to a wide geographic distribution of otherwise uniformly favorable high-input environments. Given that low-fertility P-fixing soils are widespread in SSA, broader adaptation of FyVary varieties should be examined with interested partners across SSA. Our selection in regions ranging in altitude from 900 – 1400 masl and subsequent successful testing in regions down to sea level already indicates that the range of adaptation is not limited to very specific local conditions.

## Author Contributions

MW conceived the study and planned experiments. JHC did marker assisted introgression of Pup1 into IR64 at IRRI. KK made crosses and developed populations at JIRCAS. Y.U. and D.T.L. did marker and sequence analyses. JPT, KK, SR, HNR, MFR, and MW conducted field experiments in Madagascar and collected data. MW did data analyses, prepared figures and tables and wrote the manuscript draft. MC discussed participatory plant breeding approaches and wrote part of the manuscript. All authors checked the final version of the manuscript.

## Supplementary data

**Supplementary Figure S1**: Results of hedonic test conducted at 4 villages with 100 participants each. Rice was prepared locally by villagers and their impression on appearance, taste and texture recorded by FOFIFA staff. MTM 32 and HY-85-2 correspond to FyVary32 and 85, respectively. While slight differences in preference existed between villages, with FyVary32 being preferred in Anjiro compared to FyVary85 in Marovoay, overall rankings of both varieties were not significantly different from local variety X265.

**Supplementary Figure S2**: (A) Graphical genotype of FyVary32. (B) Genotypes at the *Pup1* locus where markers K41-K59 (Chin et al. 2011) are diagnostic of gene models within the large indel missing in IR64, Nipponbare and many other modern varieties. #32 and #85 refer to VyVary 32 and 85, respectively, while #45 and #52 refer to sister lines of FyVary32 that were not released as varieties because they were phenotypically not distinct enough from #32.

