## supplementary Figure S1 for "New rice varieties with improved phosphorus-efficiency for low-input smallholder rice production in Africa"

### Slide 1
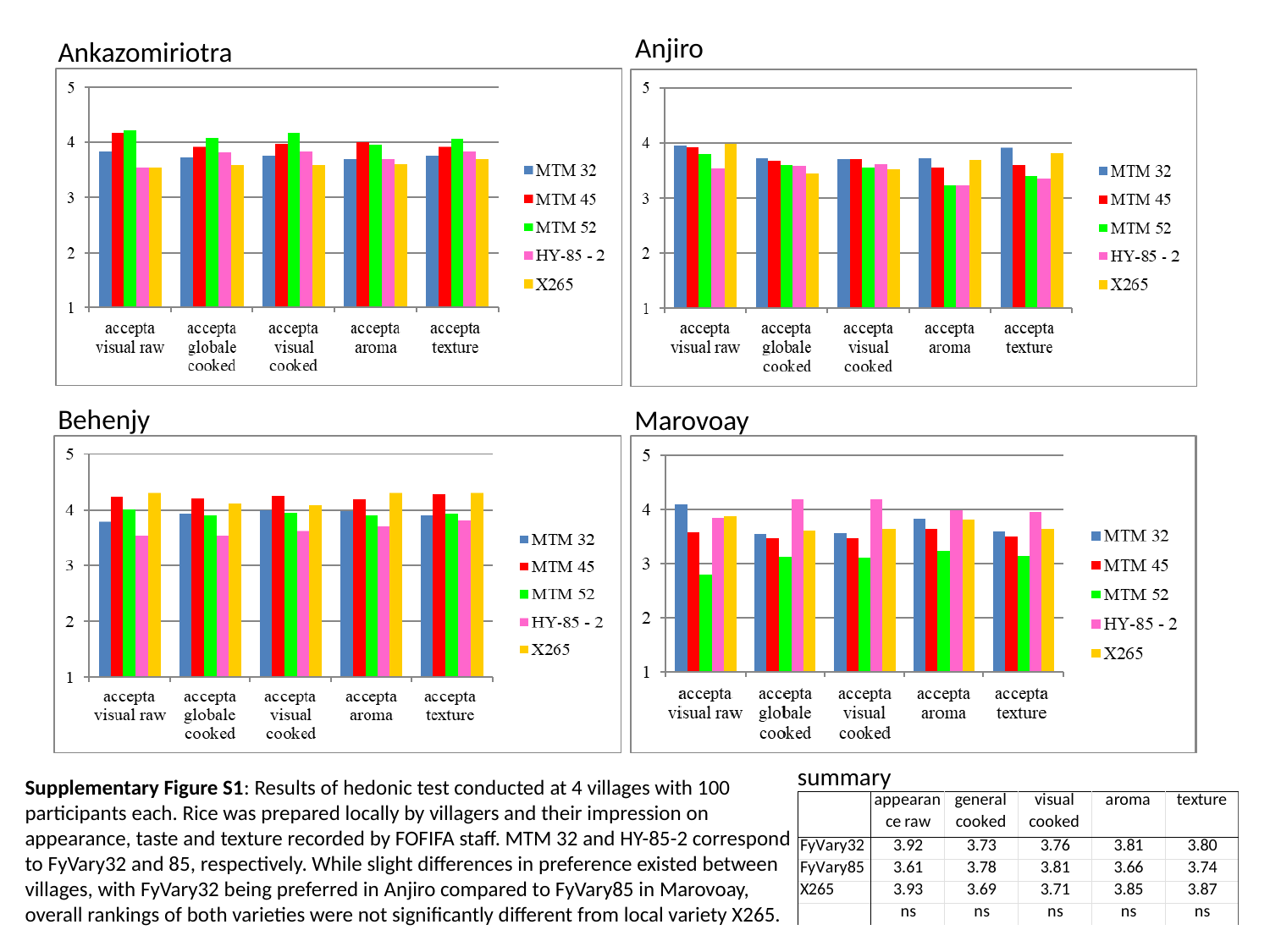

Anjiro
Ankazomiriotra
Behenjy
Marovoay
summary
Supplementary Figure S1: Results of hedonic test conducted at 4 villages with 100 participants each. Rice was prepared locally by villagers and their impression on appearance, taste and texture recorded by FOFIFA staff. MTM 32 and HY-85-2 correspond to FyVary32 and 85, respectively. While slight differences in preference existed between villages, with FyVary32 being preferred in Anjiro compared to FyVary85 in Marovoay, overall rankings of both varieties were not significantly different from local variety X265.
