## supplementary Figure S2 for "New rice varieties with improved phosphorus-efficiency for low-input smallholder rice production in Africa"

### Slide 1
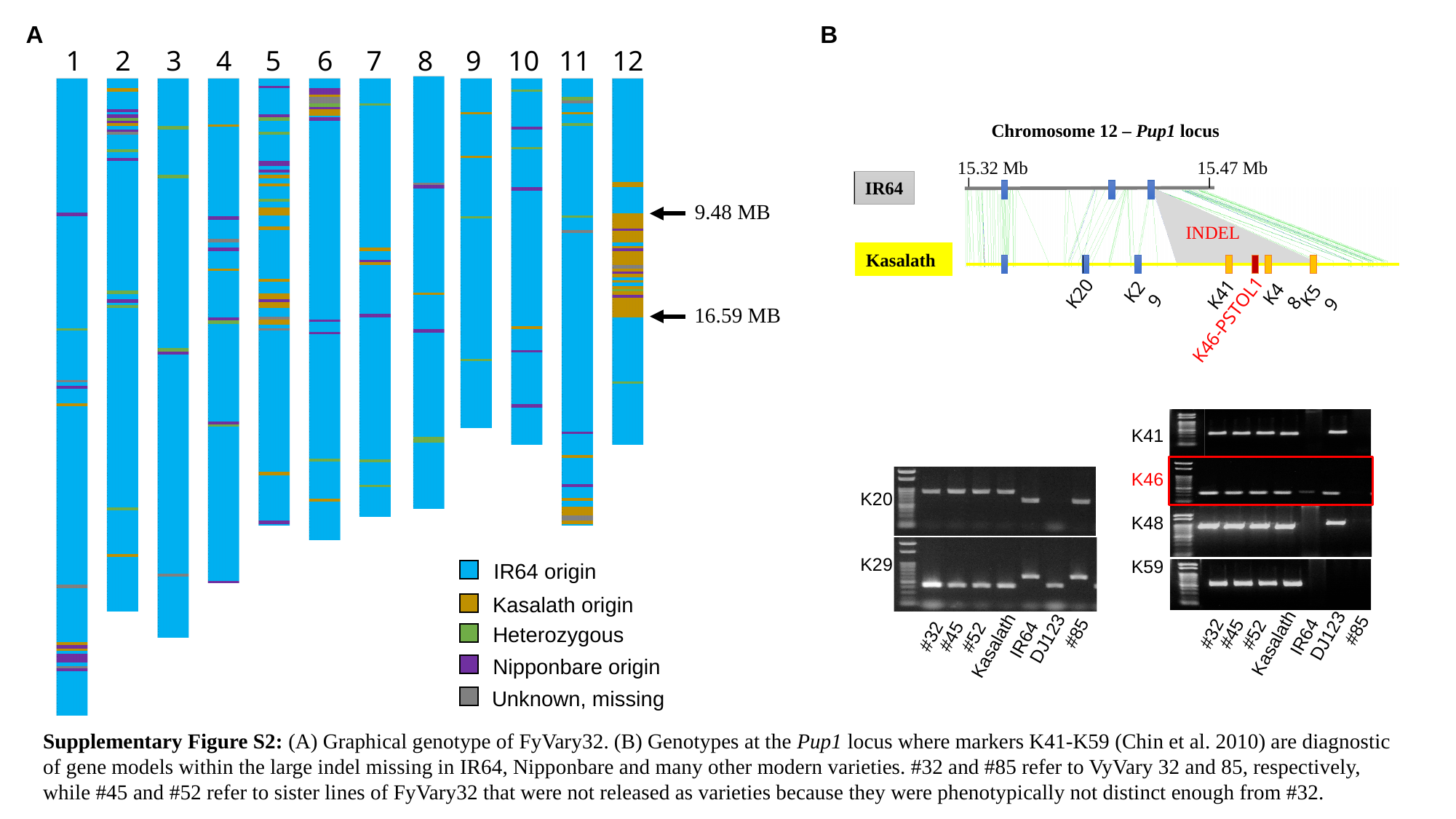

A
B
1
2
3
4
5
6
7
8
9
10
11
12
9.48 MB
16.59 MB
IR64 origin
Kasalath origin
Heterozygous
Nipponbare origin
Unknown, missing
Chromosome 12 – Pup1 locus
15.32 Mb
15.47 Mb
IR64
INDEL
Kasalath
K20
K41
K29
K48
K59
K46-PSTOL1
K41
K46
K48
K59
K20
K29
#85
#32
#45
#52
DJ123
IR64
Kasalath
#85
#32
#45
#52
DJ123
IR64
Kasalath
Supplementary Figure S2: (A) Graphical genotype of FyVary32. (B) Genotypes at the Pup1 locus where markers K41-K59 (Chin et al. 2010) are diagnostic of gene models within the large indel missing in IR64, Nipponbare and many other modern varieties. #32 and #85 refer to VyVary 32 and 85, respectively, while #45 and #52 refer to sister lines of FyVary32 that were not released as varieties because they were phenotypically not distinct enough from #32.
